# Top-down coordination of local cortical state during selective attention

**DOI:** 10.1101/2020.03.26.009365

**Authors:** Jochem van Kempen, Marc A. Gieselmann, Michael Boyd, Nicholas A. Steinmetz, Tirin Moore, Tatiana A. Engel, Alexander Thiele

## Abstract

Spontaneous fluctuations in cortical excitability influence sensory processing and behavior. These fluctuations, long known to reflect global changes in cortical state, were recently found to be modulated locally within a retinotopic map during spatially selective attention. We found that periods of vigorous (On) and faint (Off) spiking activity, the signature of cortical state fluctuations, were coordinated across brain areas along the visual hierarchy and tightly coupled to their retinotopic alignment. During top-down attention, this interareal coordination was enhanced and progressed along the reverse cortical hierarchy. The extent of local state coordination between areas was predictive of behavioral performance. Our results show that cortical state dynamics are shared across brain regions, modulated by cognitive demands and relevant for behavior.

**One Sentence Summary:** Interareal coordination of local cortical state is retinotopically precise and progresses in a reverse hierarchical manner during selective attention.

---

Cortical activity is not solely determined by external inputs but reflects ongoing fluctuations in neural excitability referred to as cortical state (Harris and Thiele, 2011; Kohn et al., 2009). Endogenous variability in cortical state shapes sensory responses and influences behavioral performance (Arieli et al., 1996; Gutnisky et al., 2017; McGinley et al., 2015a; Renart and Machens, 2014; Scholvinck et al., 2015). Although these fluctuations were long thought to be a global phenomenon that influences activity throughout the cortex (Harris and Thiele, 2011; Lee and Dan, 2012), recent evidence has revealed that signatures of cortical state are modulated locally within the retinotopic map in Macaque V4 during selective attention (Engel et al., 2016). Cortical state fluctuations manifest in periods of vigorous (On) and faint (Off) spiking activity occurring synchronously across cortical laminae. Spatially selective attention directed towards the receptive fields (RFs) of the neural population modulates On-Off dynamics by increasing the duration of On episodes (Engel et al., 2016). Thus, cognitive demands that selectively affect targeted retinotopic locations can modulate local signatures of global cortical state fluctuations. However, perception and cognition depend on activity of many areas spanning the cortical hierarchy, which begs the question of whether cortical-state dynamics are coordinated across different brain regions during attention, whether this coordination progresses in a top-down or bottom-up manner, and whether it is relevant for behavior.

We recorded simultaneously from V1 and V4 using 16-contact laminar electrodes whilst three rhesus macaques performed a feature-based spatial attention task (Fig. 1A). Electrodes were inserted perpendicular to the cortical surface on a daily basis such that RFs overlapped both across all channels within each area and between the two areas (Fig. 1B & Fig. 1C). We characterized On-Off dynamics in each area individually by fitting a Hidden Markov Model (HMM) to the spike counts (10 ms bins) of multiunit activity (MUA) across included channels (supplementary material, Fig. 1D). In line with previous reports for V4 (Engel et al., 2016), we found that a 2-phase model was the most parsimonious model for the majority of recordings (V1: 64 out of 77 recordings (83.1%), V4: 73 out of 79 recordings (92.4%), V1 and V4: 57 out of 73 recordings (78.1%), Fig. S1A-D). During these recordings, On-Off dynamics occurred without any obvious periodicity (Fig. S1E). When attention was directed towards the RFs under study, firing rates were higher during both Off and On epochs [Wilcoxon signed rank test; V1: Off *P* = 10^−164^, On *P* = 10^−87^, V4: Off *P* = 10^−184^, On *P* = 10^−103^] (Fig. 1E), On-epoch durations increased in both V1 and V4 [Wilcoxon signed rank test; V1 *P* = 10^−13^, V4 *P* = 10^−8^] and Off epoch durations increased in V1 but not V4 [Wilcoxon signed rank test; V1: *P* = 10^−8^, V4 *P* = 0.81] (Fig. 1F). Additionally, when attention was directed towards the RFs, altogether more time was spent in an On phase (Fig. S2A) and transitions to an On phase were more likely (Fig. S2B).

**Fig. 1.**
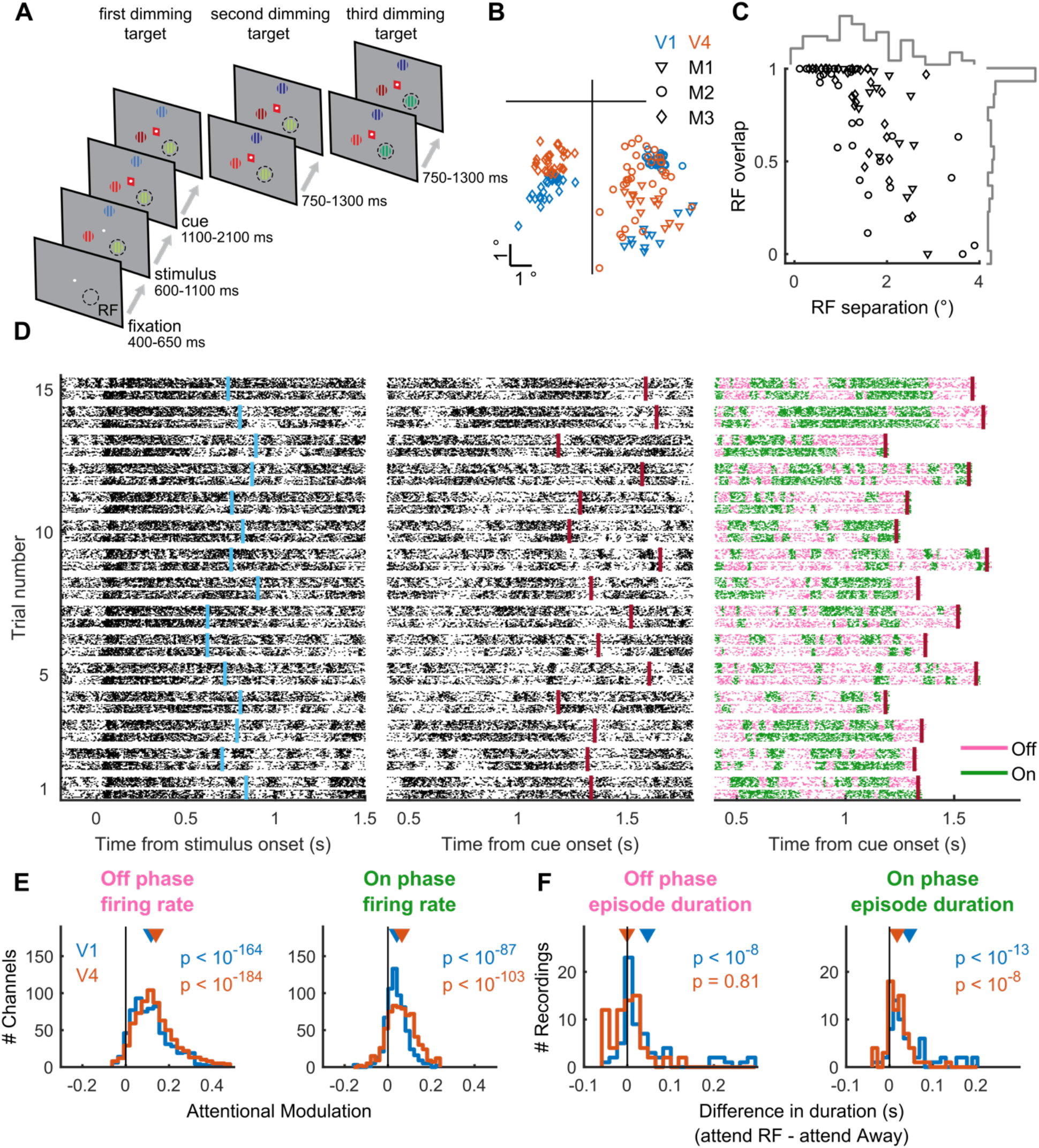
On-Off dynamics in V1 and V4 are modulated during selective attention. (**A**) Behavioral paradigm. The monkey held a lever to initiate the trial, hereafter a central fixation spot was turned on. Upon fixation 3 colored gratings appeared, one was presented inside the receptive fields (RFs) of the V1 neurons. After a variable delay a cue matching one of the grating colors surrounded the fixation spot, indicating which grating was behaviorally relevant (target). In pseudorandom order the stimuli decreased in luminance (dimmed). Upon dimming of the target, the monkey had to release the lever. (**B**) Average RF center locations (across channels) for each recording, separately for each subject (M1-M3) and area. (**C**) RF separation between V1 and V4 plotted against their overlap, expressed as the proportion of the V1 RF. The histograms along the top (right) indicate the distribution of RF separation (overlap) across all recordings. (**D**) Raster plot of HMM fit to population activity in V1 and V4. Simultaneously recorded multi-unit spiking activity on 16-contact laminar electrodes in V1 and V4 for 15 example trials, aligned to stimulus (left) and cue onset (middle and right). Each trial shows across laminar activity in V1 (bottom) and V4 (top), as raster plots (left two columns) color coded according to HMM estimation of On and Off phases (right). Middle and right columns depict the same activity. The HMM was fit from 400 ms after cue onset to 30 ms after the first dimming event. Cue onset and first-dimming are indicated for each trial by purple and red vertical bars respectively. (**E**) Attention increases firing rates during Off and On phases, both in V1 and V4. (**F**) Attention increases the duration of On episodes, both in V1 and V4, whereas it increases the duration of Off episodes only in V1. Statistics: Wilcoxon signed rank test.

We examined whether these spontaneous transitions were coordinated across visual areas. We computed cross-correlations between the V1 and V4 time series of On-Off phases (as estimated by the HMMs) during passive fixation (before stimulus onset) and during directed attention (after cue onset). During fixation, V1 and V4 transitions were coordinated but without either area leading/lagging the other [Wilcoxon signed rank test; *P* = 0.13] (Fig. 2A). During directed attention, the coordination between V1 and V4 was enhanced and On-Off transitions more often occurred in V4 before they were followed in V1, as evident from the skew towards negative values of the V4 relative to V1 transition times [Wilcoxon signed rank test; *P* < 10^−5^] (Fig. 2A). The cross-correlation strength and skew was independent of microsaccades (Fig. S3), and the strength was inversely related to the separation between V1 and V4 RFs [r = −0.38, *P* = 0.004] (Fig. 2B). Thus, the strength of On-Off dynamics coordination between visual areas is tightly coupled to their retinotopic alignment. To further characterize this interareal coordination, we computed average firing rates in V1 aligned to On-Off transition times in V4 and vice-versa. In line with transitions being driven in a top-down manner, V1 firing rate changes followed V4 transitions whereas V4 firing rate changes preceded V1 transitions (Fig. 2C). We also analyzed spiking activity simultaneously recorded with 16-contact linear electrodes inserted perpendicular to layers in V4 and tangential to layers in the frontal eye field (FEF) (or with single electrodes in FEF in some sessions) from two monkeys performing a selective attention task (V4 data reported previously in ref. (Engel et al., 2016)). A similar analysis revealed that changes of FEF firing rates precede On-Off transitions in V4 (Fig. 2D). These results suggest that On-Off transitions traverse from higher to lower areas along the visual hierarchy during selective attention.

**Fig. 2.**
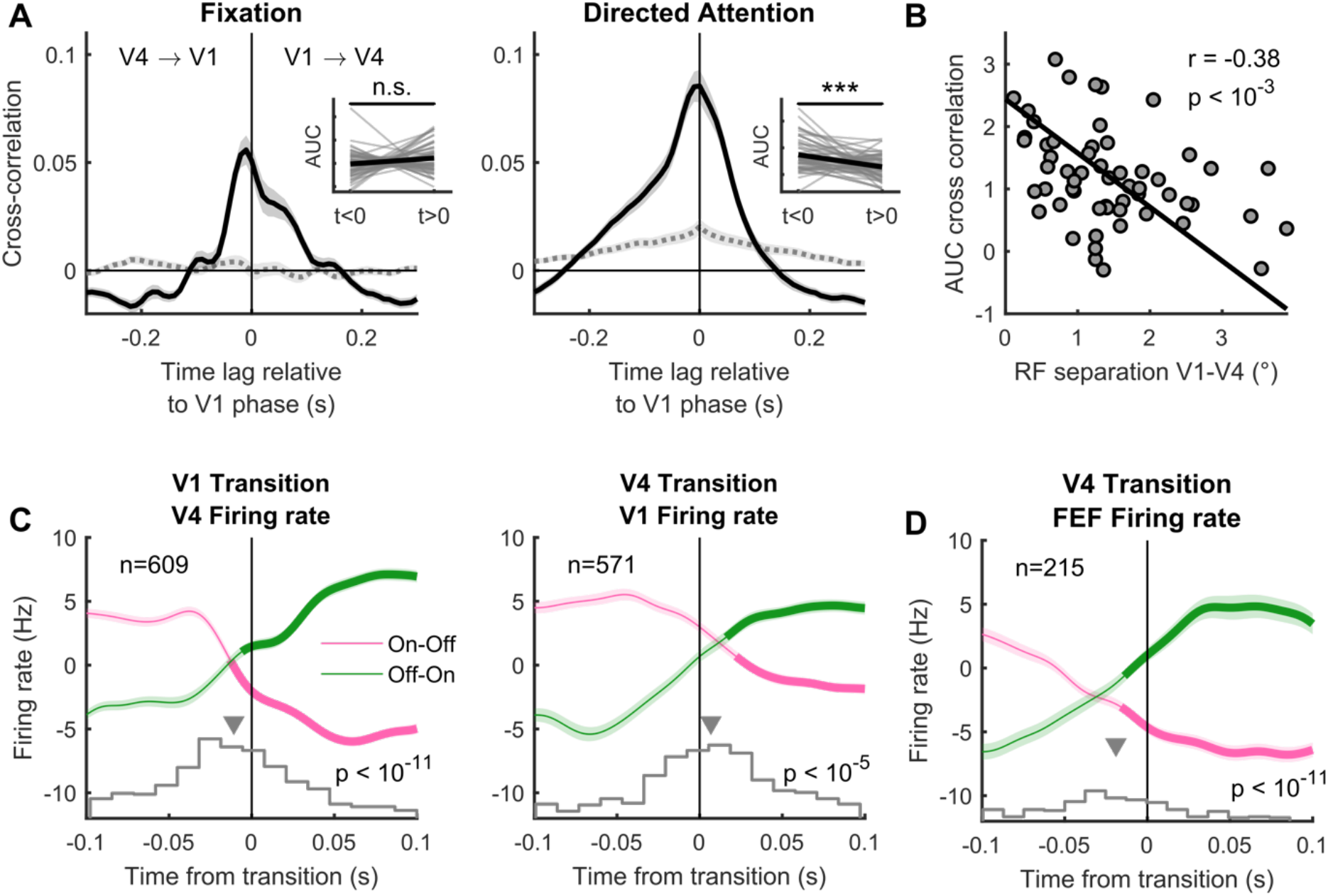
Across area coordination of cortical state. (**A**) Cross-correlation between time series of On-Off phases in V1 and V4 relative to V1 phase during passive fixation (left) and after cue onset (right). Insets show the area under the cross-correlation curve for times smaller and larger than zero. The dashed grey line depicts the shuffle predictor. (**B**) RF separation plotted against the area under the cross-correlation curve during attention (from the right panel **A**). The line indicates the standardized major axis regression fit. (**C**) Spiking activity in one area aligned to state transitions in the other area, averaged across channels and recordings. Only epochs without transitions preceding or following the alignment transition within 100 ms were included. Thick green and pink lines indicate the times the firing rate was higher (green) or lower (pink) than the average rate. Along the bottom are the histograms of the crossing point of two straight lines fit (least-squares) to the transition-aligned multi-unit firing rate. (**D**) Conventions as in **C**, but from a different dataset in which activity was recorded simultaneously from V4 and FEF. Statistics: Wilcoxon signed rank test (**A**), Pearson correlation (**B**) and FDR-corrected, one-sided, Wilcoxon signed rank test (**C** & **D**). Shaded regions denote ±1 SEM.

To investigate the relationship between V1 and V4 On-Off transitions more closely, we fit a 4-state HMM to V1 and V4 data simultaneously (HMM_V1-V4_), with the four HMM-states defined as (state 1) V1_off_-V4_off_, (state 2) V1_on_-V4_off_, (state 3) V1_off_-V4_on_ and (state 4) V1_on_-V4_on_ (Fig. 3A). This model allowed us to investigate two specific scenarios (Fig. 3B). In the first scenario (yellow), we asked: from a situation in which both areas are in an Off phase (state 1), is it more likely for V1 (state 2) or V4 (state 3) to transition (first) to an On phase? The second scenario (purple) addresses a related question: from a situation in which both areas are in an On phase (state 4), is it more likely for V1 (state 3) or V4 (state 2) to transition (first) to an Off state? The transition probabilities (Fig. 3C & Fig. 3D) revealed that when both areas were in an Off phase, it was more likely for V4 to transition to an On phase first [Wilcoxon signed rank test; *P* < 10^−3^]. Likewise, if both areas were in an On phase, it was more likely for V4 to transition to an Off phase first [Wilcoxon signed rank test; *P* < 10^−3^]. Thus, when both areas are in the same phase, it is more likely for V4 to transition away from this phase first. This finding was, however, not specific to the attend RF condition, as we found similar results for each individual attention condition (attend RF and attend away), as well as during fixation (data not shown). Selective attention, however, modulated the transition probabilities from the yellow scenario. Specifically, it decreased the probability of transitioning from state 1 to state 2, and increased the probability of transitioning from state 1 to state 3 [Wilcoxon signed rank test; *P* < 10^−2^] (Fig. 3E & Fig. 3F). Finally, this model revealed that, although On-Off phases/transitions are correlated, each area spends a substantial fraction of time in opposite phases (Fig. 3G). Selective attention specifically decreases the time spent in state 1 whereas it increases the time spent in state 3 and state 4, i.e. the states where V4 was in an On phase [Wilcoxon signed rank test; state 1 *P* < 10^−5^, state 2 *P* = 0.91, state 3 *P* < 10^−2^, state 4 *P* < 10^−3^] (Fig. 3H).

**Fig. 3.**
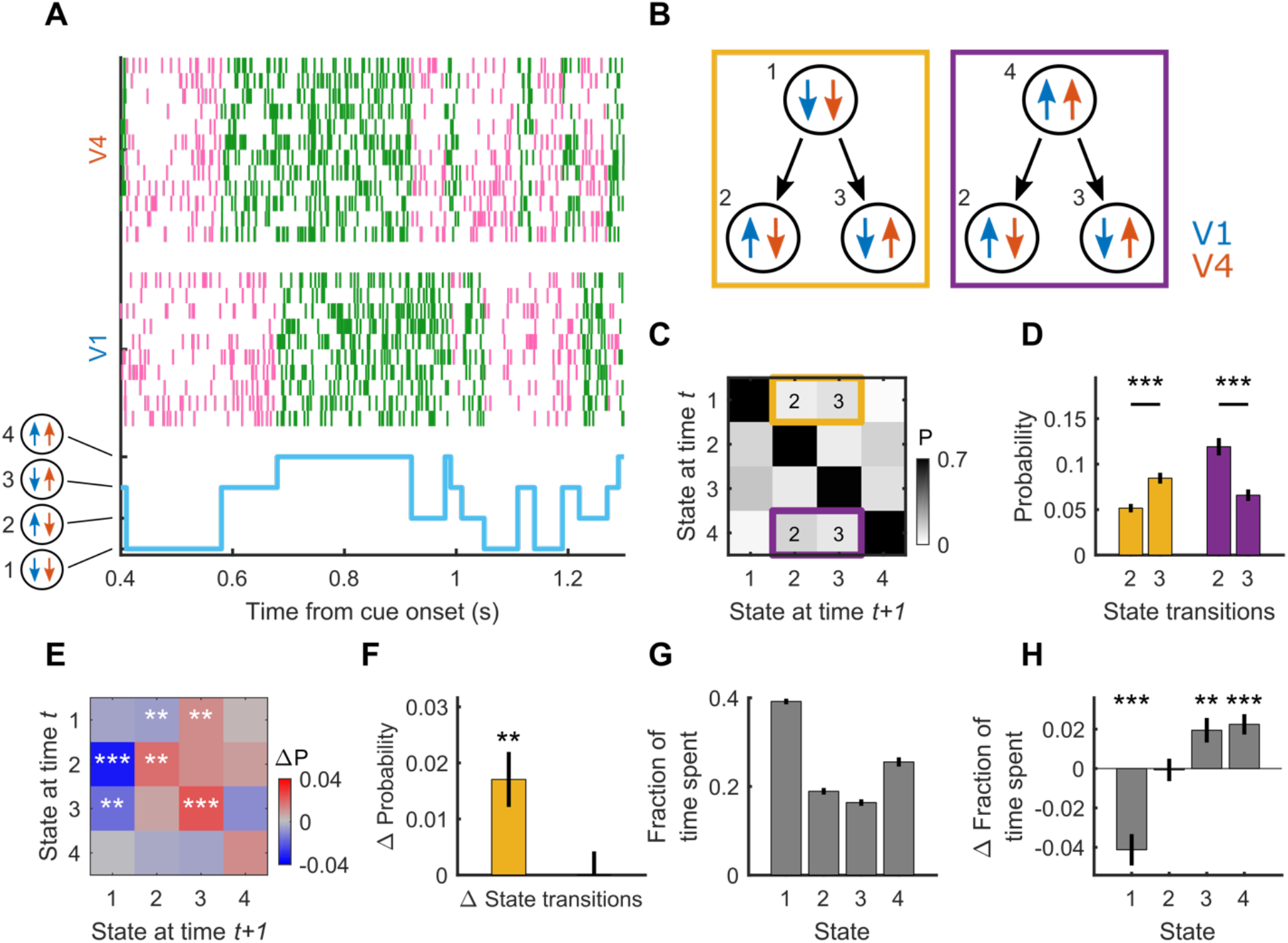
HMM with 4 states fit simultaneously to V1 and V4 data. (**A**) Example trial with the HMM state-trajectory (bottom) and across-laminar raster plot for V1 (middle) and V4 (top). (**B**) Schematic describing scenarios for testing two questions: (1, left yellow box) from a state where both V1 and V4 are Off, is it more likely for V1 or V4 to transition to the On phase first? (2, right purple box) from a state where both V1 and V4 are On, is it more likely for V1 or V4 to transition to the Off phase first? (**C**) HMM transition probability matrix, indicating the probability of staying in a state (diagonal) or transitioning from one state to another. Highlighted are scenarios set out in panel **B**. (**D**) Transition probabilities indicated in panels **B** and **C**. (**E**) Attentional influence on state-transition probabilities: shown is the difference transition matrix (attend RF – attend Away). (**F**) Attentional influence (attend RF – attend Away) on the difference between state transition probabilities (state 3 – state 2) for each of the two scenarios indicated in panels **B**, **C** and **D**. Selective attention increases the difference between the transition probabilities for state 2 and 3 for the yellow, but not the purple scenario. (**G**) The fraction of time spent in each of the 4 states. (**H**) The difference in time spent in each of the 4 states when attention is directed towards or away from the RF (attend RF – attend Away). Statistics: Wilcoxon signed rank test (FDR corrected), error bars denote ±1 SEM across recordings; *, **, *** indicate significance levels (p < 0.05, p < 0.01 and p < 0.001, respectively).

On-Off dynamics furthermore related closely to measures of (bipolar re-referenced) local field potential (LFP) (de)synchronization. During On phases in either V1 or V4, low frequency (< ^~^20 Hz) LFP power was suppressed and high frequency (> ^~^20 Hz) power was increased, both in V1 and V4 (Fig. S4A-D). Additionally, LFP power spectra in both areas varied across the 4 states of HMM_V1-V4_ (Fig. S4E-F). Here, we specifically investigated the difference in power spectra across states where the On-Off phase within an area remained constant, but differed in the other area. For example, we investigated the V1 LFP power spectra across states 1 and 3, wherein V1 was in an Off phase during both states, but V4 was either Off or On. This analysis revealed that the LFP power in V1 is modulated by V4 phase bidirectionally. If V1 was in either an On or an Off phase, a transition to an On phase in V4 increased V1 high-frequency power. A transition to an On phase in V1, however, only affected V4 high-frequency power when V4 was in an Off phase. When V4 was in an On phase, V1 phase did not affect high-frequency dynamics in V4. Thus, V4 phase influenced V1 LFP regardless of V1 phase, whereas V1 phase affected high-frequency dynamics in V4 only during Off phases in V4.

In addition to selective attention, On-Off dynamics were linked to global arousal levels, as measured by pupil diameter (Aston-Jones and Cohen, 2005; McGinley et al., 2015a, 2015b; Reimer et al., 2014; Vinck et al., 2015). For each area individually, On epoch durations were longer on trials with larger baseline pupil diameter (Fig. S5A-C), in line with previous results (Engel et al., 2016). Furthermore, pupil diameter was predictive of On-Off dynamics coordination. Larger baseline pupil diameter was predictive of shorter epoch durations for HMM_V1-V4_ state 1 (where both areas were Off) and longer state 4 epoch durations (where both areas were On) (Fig. S5D). Central arousal, in addition to focused attention, thus specifically influenced epoch durations for states where V1 and V4 phase were aligned. Importantly, pupil diameter did not differ between attention conditions (Fig. S5E), while cortical state did. This shows that effects of arousal and attention on On-Off dynamics are independently controlled.

We have demonstrated that the coordination of On-Off dynamics is retinotopically organized and driven in a top-down manner during selective attention. Is this organization also relevant for behavior? For both V1 and V4 individually, the On/Off phase at the time of target dimming was predictive of reaction time (RT) when the target was presented inside the RFs. We found an interaction between attention and On/Off phase [linear mixed effects model; V1 *β* = 0.16±0.06, *P* = 0.006, V4 *β* = 0.13±0.06, *P* = 0.03] with a main effect for phase [V1 *β* = −0.27±0.09, *P* = 0.002, V4 *β* = −0.26±0.09, *P* = 0.004], but no main effect of attention [V1 *β* = −0.15±0.09, *P* = 0.09, V4 *β* = −0.1±0.09, *P* = 0.28]. Specifically, when either area was in an On phase when the target grating dimmed, RT was faster [Wilcoxon signed rank test; V1 *P* = 0.001, V4 *P* < 10^−4^] (Fig. 4A). We furthermore found that On-Off phase coordination between V1 and V4, as assessed using HMM_V1-V4_, was also predictive of behavioral performance. Again we found an interaction between attention and On/Off phase [linear mixed effects model; *β* = 0.08±0.02, *P* < 10^−3^], with a main effect of phase [*β* = −0.15±0.04, *P* < 10^−4^], but not of attention [*β* = −0.07±0.07, *P* = 0.24]. Performance was worst when at the time of target dimming both V1 and V4 were in an Off phase (state 1). Performance improved when either area was in an On phase, and it improved even further when both areas were in an On phase at the time of target dimming (Fig. 4B). The coordination of On-Off dynamics across visual areas is thus more beneficial for behavioral performance than the state in either area alone.

**Fig. 4.**
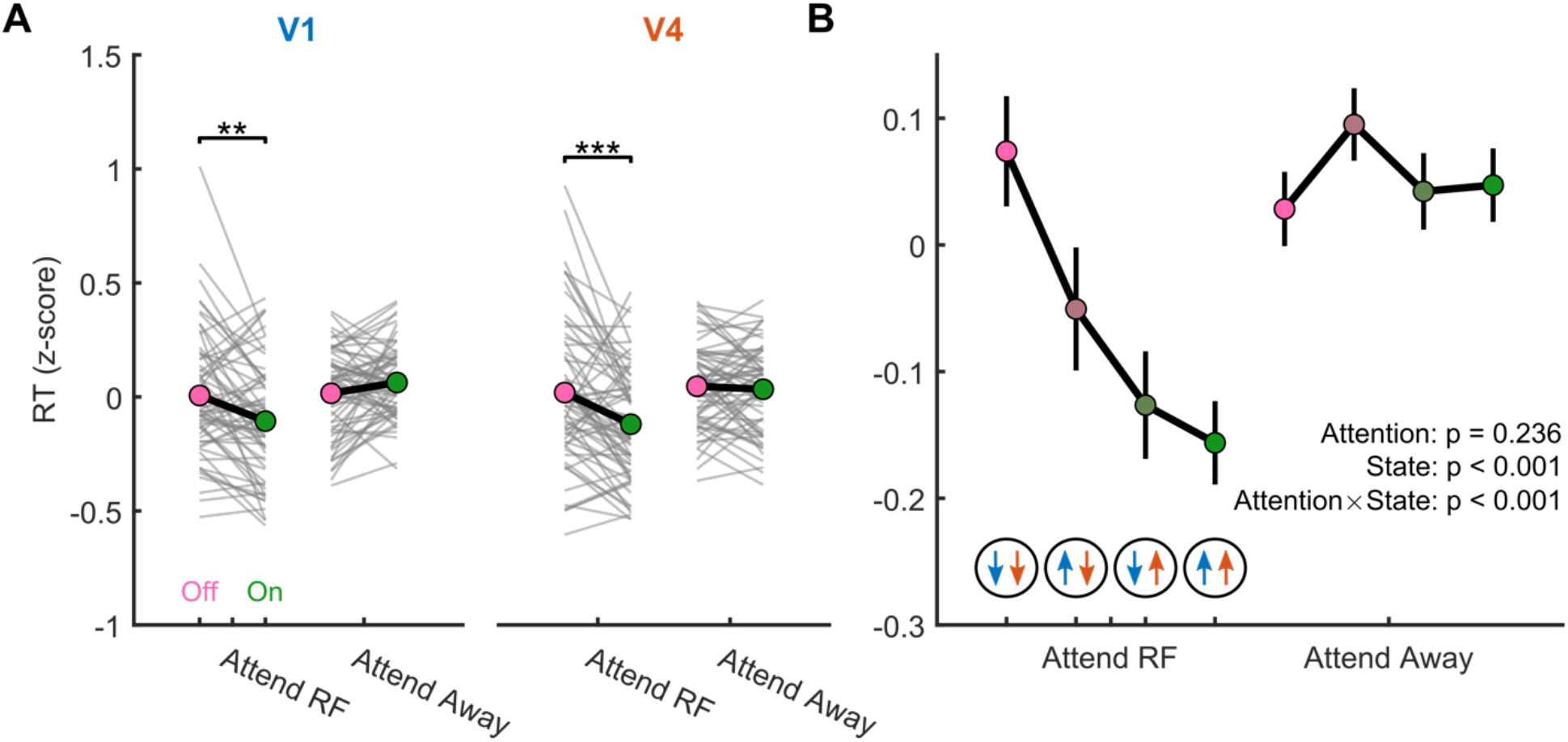
Across-area coordination of On-Off dynamics predicts behavioral performance. (**A**) On vs. Off phase of population activity at the time of target dimming, determined individually for V1 and V4, predicts behavioral performance. RT decreases when attention is directed towards the RFs and either V1 or V4 is in an On phase. (**B**) RT decreased from when both areas were Off, through V1 On - V4 Off, through V1 Off - V4 On, to V1 and V4 On when attention was directed towards the RFs. Statistics: Wilcoxon signed rank test (**A**), and multilevel linear mixed effect model (**B**). Error bars denote ±1 SEM, and *, ** and *** indicate FDR corrected significance levels of p < 0.05, p < 0.01 and p < 0.001, respectively.

In line with previous results (Engel et al., 2016), we show that transitions between On and Off phases in V4 are modulated locally by spatially selective attention. In addition, we found that they also occur in primary visual cortex (V1). Importantly, we show that interareal coordination of On-Off dynamics occurs at a local retinotopic scale, which reflects the precision of anatomical connections, and is driven in a top-down manner across areas FEF, V4 and V1 during selective attention. Fluctuations in cortical state have previously been ascribed to neuromodulatory influences (Buzsaki et al., 1988; Constantinople and Bruno, 2011; Lee and Dan, 2012) and feedback projections (Zagha et al., 2013). On-Off dynamics relate to both these mechanisms as pupil diameter, associated with neuromodulatory regulation of network state (Aston-Jones and Cohen, 2005; de Gee et al., 2017; Eldar et al., 2013; Joshi et al., 2016; Murphy et al., 2014; Reimer et al., 2016; Varazzani et al., 2015), and top-down retinotopic alignment, probably driven by feedback mechanisms (Zagha et al., 2013), are predictive of cortical state fluctuations.

The interareal coordination of On-Off dynamics and its relevance to behavioral performance suggests that trial-by-trial coordination of activity across brain regions is beneficial for information transfer and selectively modulated according to task demands. Across-area oscillatory activity is correlated according to both retinotopy and stimulus selectivity (Lewis et al., 2016). Selective attention modulates this interareal coherence (Bosman et al., 2012; Buschman and Miller, 2007; Gregoriou et al., 2009), potentially facilitating communication between hierarchically linked areas (Fries, 2005). Although attention can reduce within-area spike count correlations (Cohen and Maunsell, 2009; Herrero et al., 2013; Mitchell et al., 2009), depending on the signal correlation between neuronal pairs (Rabinowitz et al., 2015; Ruff and Cohen, 2014), it increases correlated variability across functionally related areas (Oemisch et al., 2015; Ruff and Cohen, 2016). This increased coordination might be a prerequisite for successful interareal information transfer (Harris and Mrsic-Flogel, 2013) and might allow propagation of sensory information to other brain regions (Luczak et al., 2013). When hierarchically linked areas are simultaneously active, potentially driven by the frontal cortex, global representation of information through recurrent processing could be facilitated, thereby aiding conscious stimulus processing (Baars, 2002; Dehaene and Changeux, 2011). Cognitive modulation of cortical state coordination could be a key component of this.

## Acknowledgments

We thank Demetrio Ferro for advice on data analysis.

## Author contributions

Jochem van Kempen: Conceptualization, Methodology, Formal analysis, Investigation, Data Curation, Writing – original draft preparation, Writing – review and editing, Visualization

Marc A. Gieselmann: Software, Writing – review and editing, Supervision

Michael Boyd: Investigation

Nicholas A. Steinmetz: Investigation, Writing – review and editing

Tirin Moore: Resources, Writing – review and editing, Funding acquisition

Tatiana A. Engel: Conceptualization, Methodology, Software, Writing – review and editing, Supervision

Alexander Thiele: Conceptualization, Investigation, Resources, Writing – review and editing, Supervision, Funding acquisition

## Data and materials availability

Data and materials are available upon request.

## Materials and Methods

### Animals and procedures

Subjects in our study were 3 male rhesus macaque monkeys (*Macaca mulatta*, age 10-12 years, weight 8.5-12.5 kg). All animal procedures were performed in accordance with the European Communities Council Directive RL 2010/63/EC, the National Institute of Health’s Guidelines for the Care and Use of Animals for Experimental Procedures, and the UK Animals Scientific Procedures Act. Animals were motivated to engage in the task through fluid control at levels that do not affect animal physiology and have minimal impact on psychological wellbeing (Gray et al., 2016).

### Surgical preparation

The animals were implanted with a head post and recording chambers over area V1 and V4 under sterile conditions and general anesthesia. Surgical procedures and postoperative care conditions have been described in detail previously (Thiele et al., 2006).

### Behavioral paradigm

Stimulus presentation and behavioral control was regulated by Remote Cortex 5.95 (Laboratory of Neuropsychology, National Institute for Mental Health, Bethesda, MD). Stimuli were presented on a cathode ray tube (CRT) monitor at 120 Hz, 1280 × 1024 pixels, at a distance of 54 cm. The location and size of receptive field (RF) were measured as described previously (Gieselmann and Thiele, 2008), using a reverse correlation method. Briefly, during fixation, a series of black squares (0.5-2° size, 100% contrast) were presented for 100 ms at pseudorandom locations on a 9 × 12 grid (5-25 repetitions for each location) on a bright background. RF eccentricity ranged from 3.4 - 7.5° in V1, and from 2.5 to 8.9° in V4.

During the main task (Fig. 1A), the monkeys initiated a trial by holding a lever and fixating on a central white fixation spot (0.1°) displayed on a gray background (1.41 cd/m^2^). After a fixed delay (614, 424, 674 ms, for monkeys 1, 2 and 3), three colored (for color values see Table S1) square wave gratings appeared equidistant from the fixation spot, one was centered on the RF of the V1 neurons under study. The locations of colored gratings were fixed for each recording session but were pseudorandomly assigned across sessions. Stimulus size varied between 2 and 4° diameter, depending on RF eccentricity and size. For most recordings we used drifting gratings but presented one monkey with stationary gratings during 22 out of 34 recording days. The drifting gratings moved perpendicular to the grating orientation, with the motion direction pseudorandomly assigned on every trial. After a random delay (618-1131 ms for monkey 1, 618-948 ms for monkeys 2 and 3; uniformly distributed), a central cue appeared that matched the color of one of the gratings, indicating that this grating would be behaviorally relevant on the current trial. After a variable delay (1162-2133 ms for monkey 1, 1162-1822 ms for monkeys 2 and 3; uniformly distributed), one of the pseudorandomly selected gratings changed luminance (for color values see Table S1), referred to as dimming. If the cued grating (target) dimmed, the monkey had to release the lever in order to obtain a reward. If, however, a non-cued grating (distractor) dimmed, the monkey had to ignore this and keep hold of the lever until the target dimmed on the second or third dimming event (each after another 792-1331 ms for monkey 1; 792-1164 ms for monkeys 2 and 3; uniformly distributed).

### Data acquisition and analysis

We recorded from all cortical layers of visual areas V1 and V4 using 16-contact laminar electrodes (150 μm contact spacing, Atlas silicon probes). Out of a total of 77 V1 and 79 V4 recording sessions, 73 recordings were conducted simultaneously in both areas. The electrodes were inserted perpendicular to the cortex on a daily basis.

Raw data were collected using Remote Cortex 5.95 and by Cheetah data acquisition interlinked with Remote Cortex 5.95. Neuronal data were acquired with Neuralynx preamplifiers and a Neuralynx Digital Lynx amplifier. Unfiltered data were sampled with 24 bit at 32.7 kHz and stored to disc. Data were replayed offline, sampled with 16-bit and band-pass filtered at 0.5-300 Hz and down sampled to 1 kHz for local field potential (LFP) data, and filtered at 0.6-9 kHz for spike extraction. Eye position and pupil diameter was recorded at 220 Hz (ViewPoint, Arrington Research). Pupil diameter was recorded for 75 (90.4 %) of recordings.

All data analyses were performed using custom written Matlab (the Mathworks) scripts.

### Data preprocessing, trial selection and channel selection

We corrected for any noise common to all channels via common average reference, in which the average of all channels is subtracted from each individual channel. We extracted population activity by progressively lowering spike extraction thresholds until approximately 100 Hz spiking activity was detected on each channel between fixation onset and the first dimming event. Next, we computed the envelope of MUA (MUAe) by low-pass filtering (<300 Hz, fifth order Butterworth) the rectified 0.6-9 kHz filtered signal. Because we noticed that during some recording sessions the electrode seemed to have moved (e.g. due to movement of the monkey), we visually inspected the stability of each recording by investigating the stimulus aligned firing rates, MUAe and their baseline (−500 to −50 ms) energy across all trials and channels. With energy (*E*) defined as:

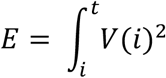

where *t* is the number of time points in the vector (*V*) representing the single-trial histogram or MUAe. We selected the largest continuous time window that showed stable activity across all V1 & V4 channels.

In addition to selecting trials from stable periods, we selected channels for further processing that were determined to be in gray matter. Using current source density (CSD), we investigated on which channels currents were entering (sinks) and exiting (sources) cortical tissue, which allowed us to determine the relative recording depth compared to the known cortical anatomy (Schroeder, 1998; Schroeder et al., 1991). The CSD profile can be calculated according to the finite difference approximation, taking the inverse of the second spatial derivative of the stimulus-evoked voltage potential *φ*, defined by:

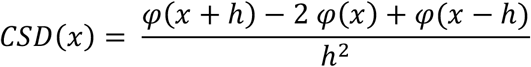

where *x* is the depth at which the CSD is calculated and *h* the electrode spacing (150 μm). We used the iCSD toolbox (Pettersen et al., 2006) to compute the CSD. With this toolbox we used a spline fitting method to interpolate *φ* smoothly between electrode contacts. We used a diameter of cortical columns of 500 μm (Mountcastle, 1957), and tissue conductance of 0.4 Sm^−1^ (Logothetis et al., 2007).

To aid determination of recording depth, we computed the signal-to-noise ratio (SNR), the response latencies to stimulus onset for each channel and the receptive field (RF) estimation (see below). SNR was computed as:

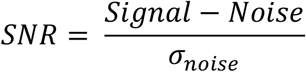

with *Signal* defined as the average MUAe amplitude in one of eight 50 ms time windows, from 30 to 80 ms, in 10 ms steps, to 100 to 150 ms after stimulus onset, and *Noise* as the average MUAe amplitude during the baseline period (−200 to 50 ms) before stimulus onset. SNR in at least one of these eight estimates was required to be higher than 3 for a channel to be included for further analyses.

We computed the response latency to stimulus onset for each channel according to the method described by Roelfsema et al. (2007). We fitted the visual response as a combination of an exponentially modified Gaussian and a cumulative Gaussian using a non-linear least-squares fitting procedure (function lsqcurvefit) to the average MUAe time course. There are two assumptions implicit in this method. First, the onset latency has a Gaussian distribution across trials and across neurons that contribute to the MUAe, and second, that (part of) the response dissipates exponentially. The visual response *y* across time *t* was modelled as:

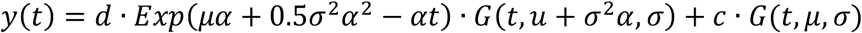

where *μ* is the mean, *σ* is the standard deviation, *α*^−1^ is the time constant of the dissipation, *G*(*t, μ, σ*) is a cumulative Gaussian, and *c* and *d* are the factors scaling the non-dissipating and dissipating modulation of the visual response. The response latency was defined as the time point where *y*(*t*) reached 33% of the maximum of the earliest peak, the first Gaussian (Roelfsema et al., 2007; Self et al., 2013). Data were aligned to the earliest current sink, the presumed thalamic input layer (L4); channels were excluded if they were >1 mm more superficial or >0.75 mm deeper than this layer.

### Receptive field estimation

Offline RFs were determined for each channel via reverse correlation of the MUAe signal to stimuli (0.5 – 2 ° black squares) presented on a 9 × 12 grid (Gieselmann and Thiele, 2008). The stimulus-response map was converted to z-scores, after which the RF for each channel was indicated by a contour (thresholded at a z-score of 3) surrounding the peak activity. These z-scored maps were averaged across all channels for each area (the population average z-score was computed using Stouffer’s Z-score method according to 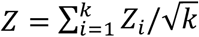, with *k* as the number of channels, after which we determined the overlap and separation between the V1 and V4 RFs (Fig 1B-C).

### Bipolar re-referencing

To ensure that global signals, common to multiple channels, did not affect our LFP and spectral analyses (see below), we re-referenced our LFP signals according to the bipolar derivation. Bipolar re-referenced LFP signals (LFPb) were computed by taking the difference between two neighboring channels.

### Attentional modulation

The effect of selective attention on neural activity was computed via an attention modulation index (*attMI*), defined as:

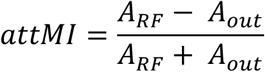

with *A_RF_* as the neural activity when attention was directed towards the RF, and *A_out_* the activity when attention was directed away from the RF. This index ranges from −1 to 1, with zero indicating no attentional modulation and with positive (negative) values indicating higher (lower) activity when attention was directed towards the RF.

### Hidden Markov Model

To quantify On-Off dynamics in V1 and V4, we fit a Hidden Markov Model (HMM) to the population activity across all laminae. We fit the HMM both to activity from each individual area, following the procedures described by Engel et al. (2016), as well as to the activity from both areas simultaneously.

Our HMM assumes that spike counts on the recorded channels can be well characterized as a doubly-stochastic process, of which the parameters can be accurately estimated (Rabiner, 1989). In this study, spike counts on each channel are assumed to be produced by a Poisson process with different (constant) mean rates during On or Off phases of the underlying ‘hidden’ (latent) process *s* common to all channels that we need to infer (Engel et al., 2016). The mean firing rate on each channel *j* in phase *s* is defined by entry 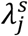 in the emission matrix Λ. The transition matrix *P* gives the probabilities of transitioning between these latent phases. In the transition matrix *P*, each entry indicates the probability of transitioning between two specific phases. For instance, *P*_11_ indicates the probability of transitioning from *s* = 0 to *s* = 0 (remaining in the Off phase), whereas *P*_12_ indicates the probability of transitioning from *s* =0 to *s* = 1, more formally: *P_11_* = *P_off_* = *P*(*s*_*t*+1_ = 0|*s_t_* = 0), *P*_12_ = 1 − *P_off_* = *P*(*s*_*t*+1_ = 1|*s_t_* = 0). These probabilities do not depend on time: at any time step *t*, the probability of transitioning between phases depends only on the value of *s* at time *t*(*s_t_*). The latent dynamics estimated by the HMM thus follow a discrete time series in which *s_t_* summarises all information before time *t*. For each channel, MUA was discretized by determining spike counts in 10 ms bins following each time *t*, with the probability of observing spike count *n* on channel *j* during phase *s* defined as

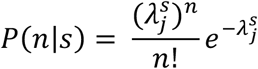

The full description of an HMM is given by the emission matrix Λ, transition matrix *P* and the probabilities *π*^0^ that indicate the initial values *s*_0_, in which 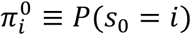. These parameters were estimated using the Expectation Maximization (EM) algorithm (Bishop, 2006), maximizing the probability of observing the data given the model according to the Baum-Welch algorithm (Rabiner, 1989). Because the EM procedure can converge to a local maximum, rather than the global maximum, we repeated the EM procedure ten times with random parameter initializations, and chose the model with the highest likelihood. Random values were drawn from Dirichlet distributions for *π*^0^ and P, and from a uniform distribution between zero and twice the channel’s mean firing rate for Λ. The EM procedure was terminated if the relative change, computed as |*new – original*|/|*original*|, in the log-likelihood was smaller than 10^−3^ and the change in the transition and emission matrix was smaller than 10^−5^, or if it reached the maximum number of iterations (n = 500).

Once the optimal parameters were estimated, we used the Viterbi algorithm to determine the most likely latent trajectory for each individual trial. We applied the HMM separately to each attention condition. For every trial, we applied the HMM during multiple time periods of the task, during fixation and during the time window from 400 ms after cue onset to 30 ms after the first dimming event. For the behavioral analysis, we additionally analyzed the period up to 30 ms after the second dimming event for trials in which target dimming did not occur on the first dimming event, and for which the first distractor dimming was not inside the RFs.

To determine what number of latent phases best described the data, we fit HMMs with the number of phases ranging from 1 to 8, and used a four-fold cross-validation procedure to compute the leave-one-channel-out cross-validation error for each HMM (Engel et al., 2016). We fit the HMM to a randomly selected subset of 3/4 of the trials, and computed the cross-validation error on the remaining 1/4 of trials. This procedure was repeated 4 times using a different 3/4 of trials for training and 1/4 of trials for testing the HMM. We computed the cross-validation error *CV_var_* for each channel *j* across all trials *K* and time bins *T* as the difference between the actual and expected spike count according to:

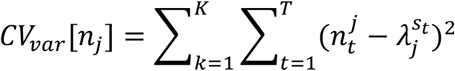

We normalized *CV_var_* to the error in the 1-phase HMM, averaged across channels, cross-validations and conditions, and determined the difference in *CV_var_* with each additional phase in the HMM. The normalized mean cross-validation error across each of the eight HMM models for all recordings is depicted in Fig. S1. For most recordings, and for both V1 and V4, *CV_var_* decreased with the addition of a second phase, but did not decrease much further with additional phases. This allowed the identification of the elbow (kink) in this error plot as the model with two phases. We included areas/recordings for further analysis that revealed a reduction in cross-validation error of at least 10% with the addition of a second phase, but did not decrease by more than 10% with additional phases. For a small subset of recordings, a three or a four-phase model fit the data best; these recordings were excluded from further analysis. In total, we found a reduction of >10% in cross-validation error when fitting a 2-phase versus 1-phase model in 64 V1 (83.1 %), and 73 V4 (92.4 %) recordings; in 57 (78.1 %) recordings we found evidence for a 2-phase model in both V1 and V4 (Fig. S1).

To investigate the across-area coordination of On-Off dynamics, we fit a 4-state HMM to V1 and V4 data simultaneously. Across these four states, both V1 and V4 could be in either an Off or On phase, with the states defined as: *V*1_*off*_ – *V*4_*off*_ (state 1), *V*1_*on*_ – *V*4_*off*_ (state 2), *V*1_*off*_ – *V*4_*on*_ (state 3) and *V*1_*on*_ – *V*4_*on*_ (state 4). This model was fit according to the same steps as the HMM applied to individual areas, with one exception. For each channel *j*, the emission rate *λ* was constrained to be the same across states for which this channel (area) was in the same phase. For example, rates were constrained for a V1 channel across state 1 and state 3, during which V1 was in an Off phase 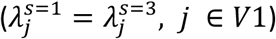.

### Testing the effect of On-Off dynamics on behavioral performance

To determine the effect of On-Off dynamics and their across-area coordination on behavioral performance, we investigated whether the On/Off phase of population activity at the time of target dimming influenced reaction times (RT). To this end, we averaged, for each recording, the RT across all trials that ended in the same phase. We subsequently tested for a relationship between On/Off phase and RT across recordings (Statistical testing).

### Cross correlation

The temporal relationship between On-Off time series and transitions, microsaccade onset times and activity in V1, V4 and FEF were investigated using cross-correlations. The cross-correlations based on HMM time series (*CC_HMM_*) were calculated using the function xcorr in Matlab, according to:

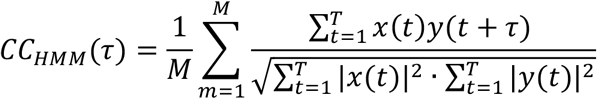

where M is the number of trials, *T* is the number of discrete time bins, *x* and *y* the mean subtracted On-Off time series in V1 and V4 as determined by the HMM, and *τ* the time lag. Here, the numerator indicates the cross-covariance, which is normalized (the denominator) such that the autocorrelation for each time series at zero lag is 1. This procedure normalized *CC_HMM_* such that correlation coefficients were obtained. We furthermore subtracted the shuffle predictor *CC_shuffle_* from *CC_HMM_* to remove any task-related (event-locked) correlations between *x* and *y*. *CC_shuffle_* was computed by shuffling *y* trials.

Cross-correlations (*CC*) between state transitions and microsaccade onset times were computed in the same way but for a different normalization (denominator) factor. Here we normalized by the number of microsaccades, resulting *CC* to be of the order of coincidences of state transitions per microsaccade.

To investigate the neural activity around the time of On-Off transitions, we computed the transition-triggered average (TTA). The TTA was estimated by computing the cross covariance (the numerator), divided by the number of transitions for each channel (denominator). Again, we subtracted the shuffle predictor to remove any task-related correlations.

### Power estimation

We estimated the power spectra of the LFPb using a custom multitaper approach based on the Chronux toolbox (Bokil et al., 2010). We estimated the power separately for On and Off states determined by the HMM using only epochs that lasted longer than 250 ms. Because epoch durations were variable, we zero-padded each segment to the next highest power of 2 of the longest epoch duration (2048 time points), ensuring we could extract the same frequencies for each segment. This approach gave us a half bandwidth (*W*) of approximately 1.95 Hz, according to *W* = (*K* + 1)/2*T*, with *K* being the number of data tapers (*K* = 7) and *T* the length of the time window in seconds. Frequencies were estimated from 4 to 200 Hz.

### Microsaccade detection

We low-pass filtered the horizontal and vertical eye traces at 30 Hz (2^nd^ order Butterworth filter) after which we detected microsaccades by using the algorithm developed by Engbert and Kliegl (2003). This algorithm converts eye position to velocity and classifies an eye movement as a microsaccade if the velocity is larger than a threshold for at least three consecutive time points. The threshold is set to 6 times the median estimator, given by: 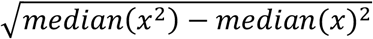, where *x* is the eye position channel. Thus, the threshold is determined for each single trial. The use of the median estimator ensured that microsaccade detection is relatively robust to different levels of noise.

### Statistical testing

To determine whether there were significant differences between attention conditions or HMM states (e.g. in firing rate or epoch duration) we made use of multiple statistical methods. We used (paired-sample) Wilcoxon signed rank tests whenever a comparison was made between two conditions (e.g. attend RF versus attend away), or to test whether a distribution was significantly different from zero. When a comparison involved multiple conditions, or multiple factors (e.g. attention and state), we used linear mixed effect models to test for main effects of each condition/factor and interaction effects between factors. These factors were defined as fixed effects and we included random intercepts for each recording as random effects, accounting for the repeated measurements within each recording. Specifically, we modelled RT as a linear combination of attention condition (*Att*) and HMM state coefficients, as well as their interaction:

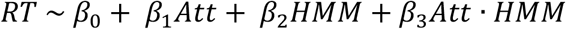

We used false discovery rate (FDR) to correct for multiple comparisons (Benjamini and Yekutieli, 2001). Error bars in all figures indicate the standard error of the mean (SEM).

**Fig. S1.**
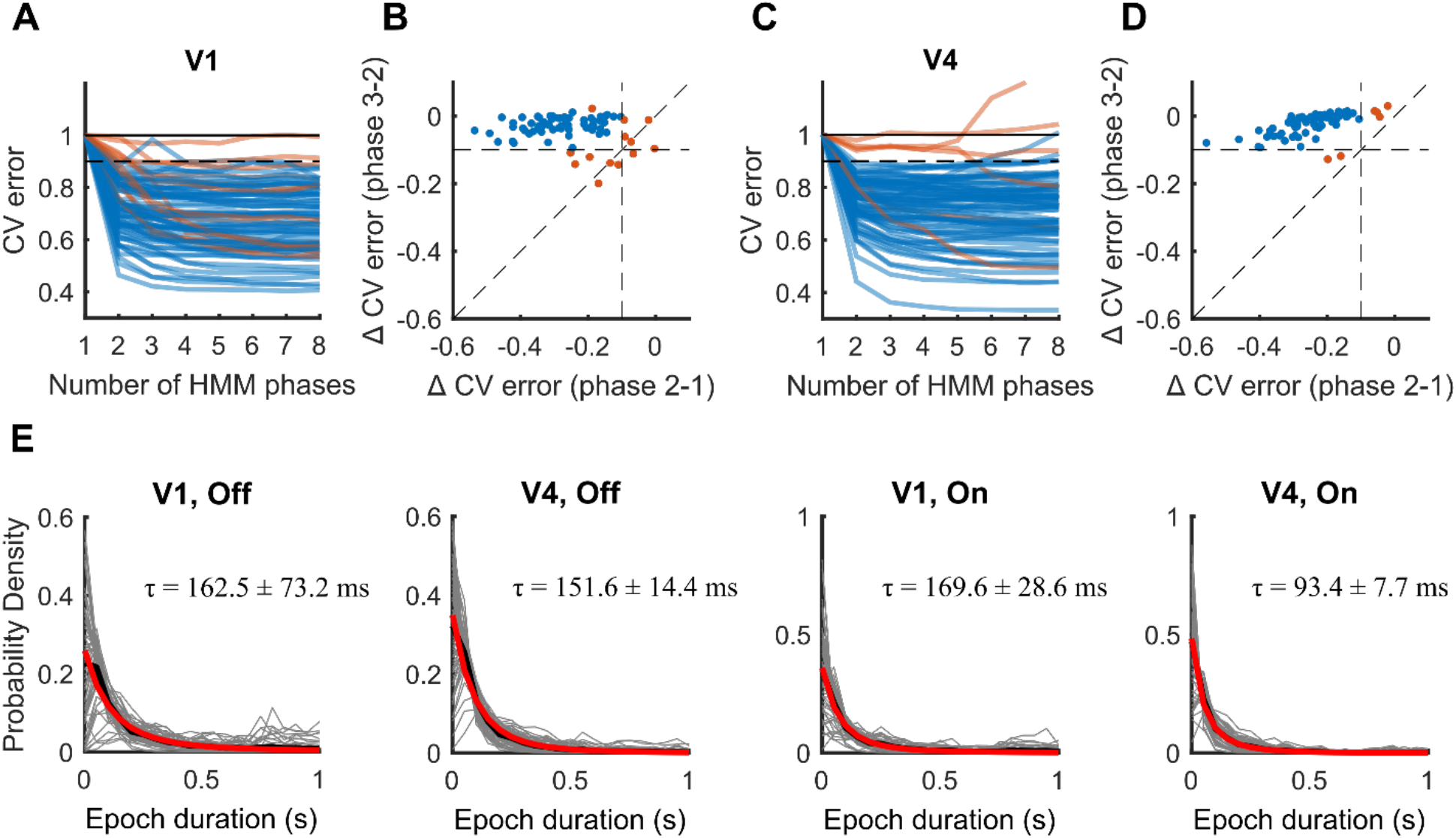
Determining the number of HMM phases and their epoch durations in V1 and V4 MUA. (**A**) Cross validation (CV) error plotted against the number of phases in each HMM for V1. (**B**) The difference in cross validation error between the 1-phase and 2-phase model, plotted against the difference between the 2-phase and 3-phase model for V1. Most recordings show a large reduction in cross-validation error with the addition of a second phase, and only marginal changes with additional phases. Blue (red) lines and markers indicate the recordings included (excluded) for further analysis. (**C-D**) Same conventions as (**A-B**) but for V4. (**E**) Distributions of Off and On episode durations overlaid by the exponential distributions with the decay constant set by the HMM transition probabilities (red, *N*(*t*) = *N*_0_*e^−t/τ^*, where *N*_0_ is the normalization constant, and *τ* is the decay time-constant computed for each recording and phase). A good match for these models indicates that On-Off dynamics were not driven by an oscillatory phenomenon. Grey and thick black lines indicate individual recordings and their mean, respectively.

**Fig. S2.**
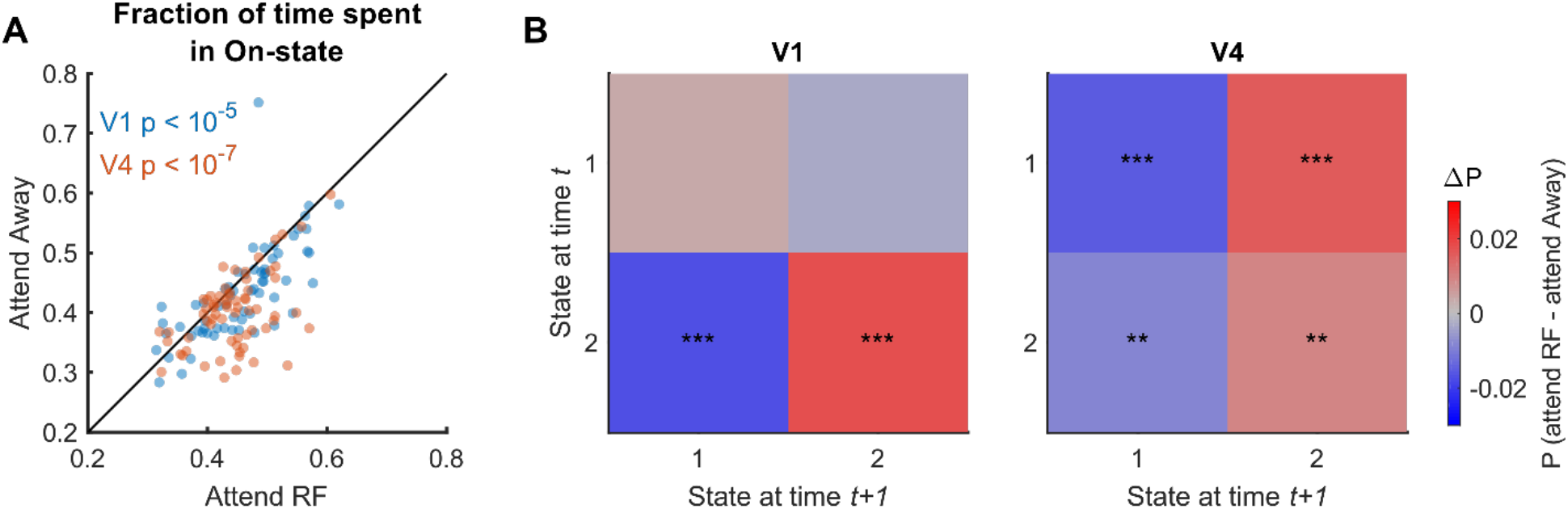
Attentional modulation of HMM parameters. (**A**) The fraction of time spent in an On phase is increased when attention is directed towards the RFs. (**B**) Attentional influence on HMM transition probabilities. Shown is the difference between transition matrices (attend RF – attend Away). Statistics: Wilcoxon signed rank tests; *, **, ***, indicate FDR corrected significance levels of p < 0.05, p < 0.01 and p < 0.001, respectively.

**Fig. S3.**
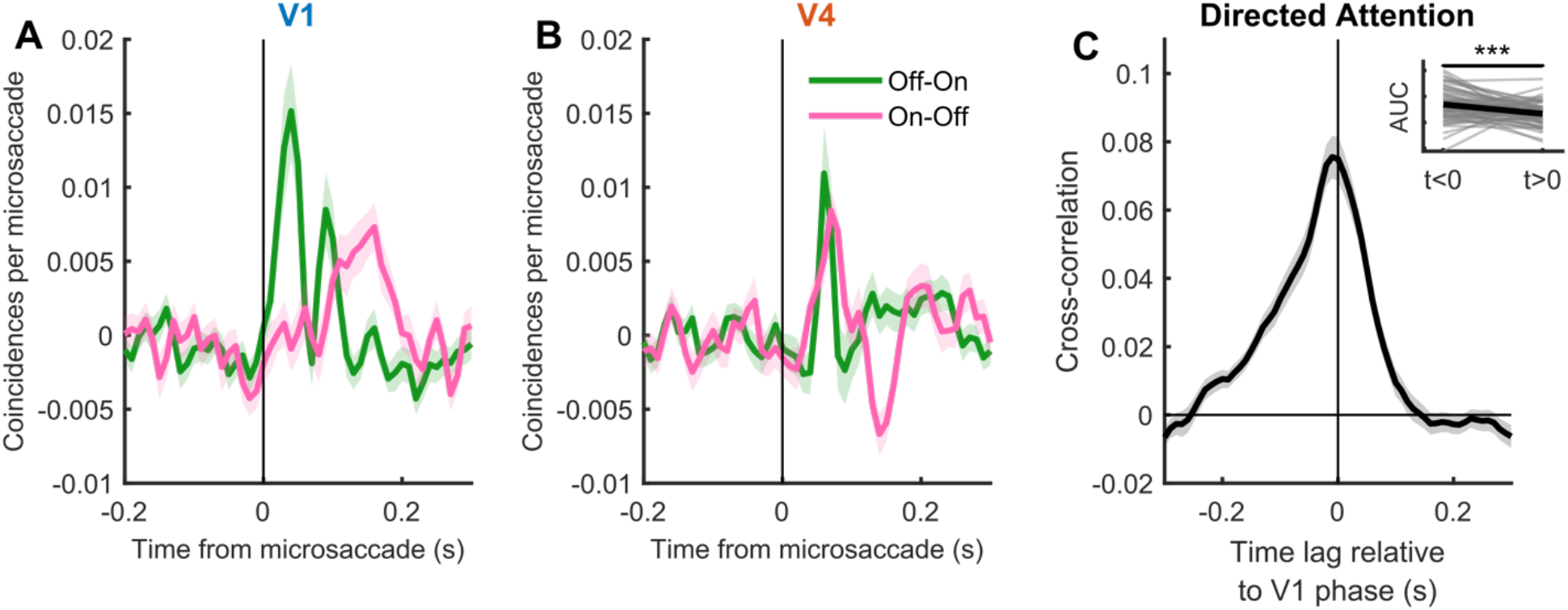
Relationship between microsaccades and On-Off transitions. (**A**) Cross-correlation of On-Off transitions in V1 triggered to microsaccade onset. (**B**) Same as panel **A**, but for On-Off transitions in V4. (**C**) Cross-correlation between time series of On-Off dynamics in V1 and V4 after exclusion of trials in which microsaccades occurred. Statistics: Wilcoxon signed rank test. Shaded regions denote ±1 SEM across recordings, *, ** and *** indicate significance levels of p < 0.05, p < 0.01 and p < 0.001 respectively.

**Fig. S4.**
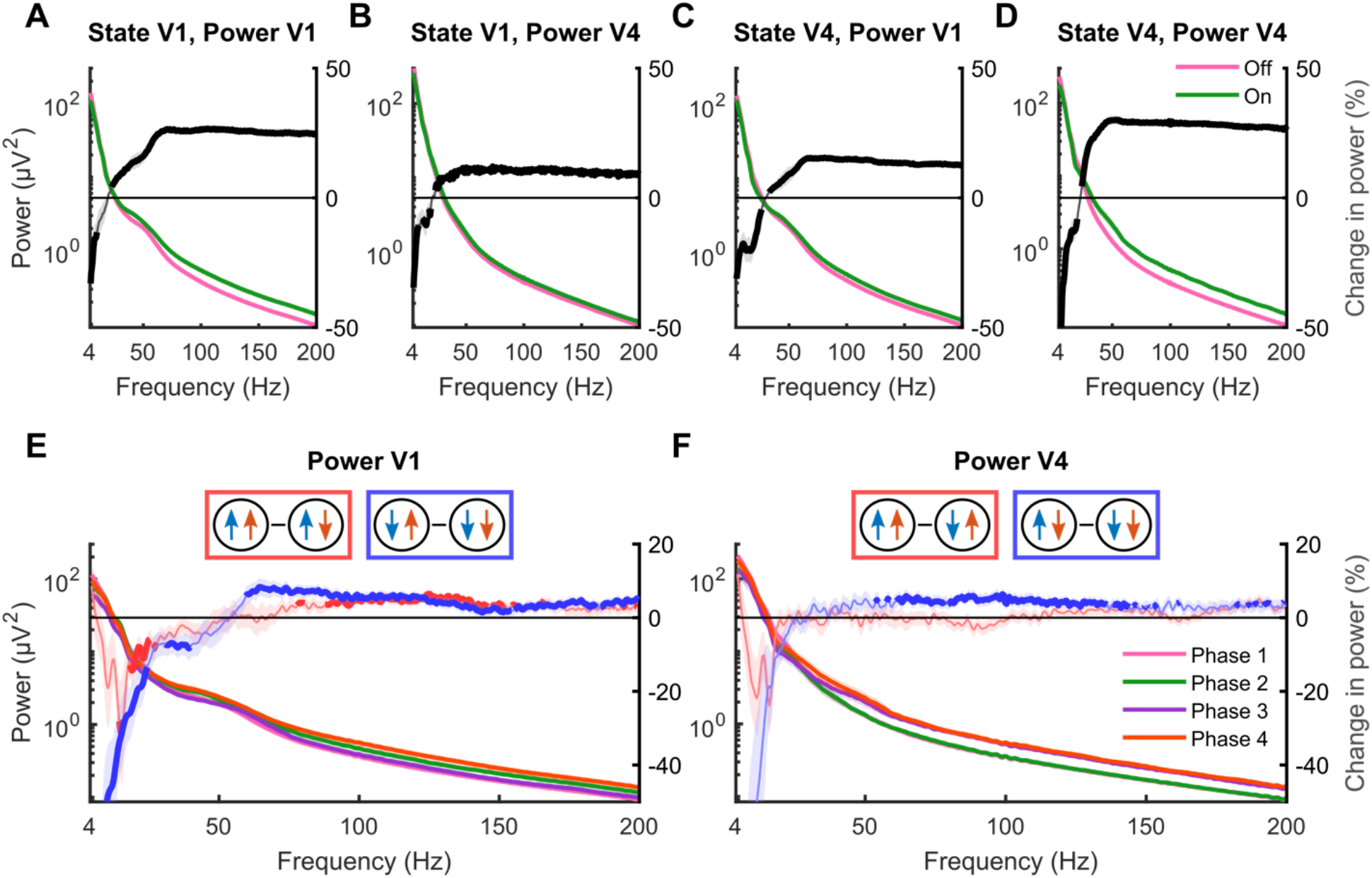
Bipolar re-referenced LFP power spectrum across HMM states. (**A**) Power in V1 during On and Off phases in V1. (**B**) Power in V4 during On and Off phases in V1. (**C**) Power in V1 during On and Off phases in V4. (**D**) Power in V4 during On and Off phases in V4. Right y-axis indicates the percentage change in power during On versus Off phases (On-Off). (**E-F**) Power spectrum in V1 (**E**) and V4 (**F**) for the 4-state HMM fit across V1 and V4 and the within-area power difference between On phases (red, V1: state 4-2; V4: state 4-3), or Off phases (blue, V1: state 3-1; V4: state 2-1). Only On/Off episodes of at least 250 ms were included. Thick percentage change lines indicate significantly modulated frequencies (p < 0.05, Wilcoxon signed rank test, FDR corrected). Shaded regions denote ±1 SEM.

**Fig. S5.**
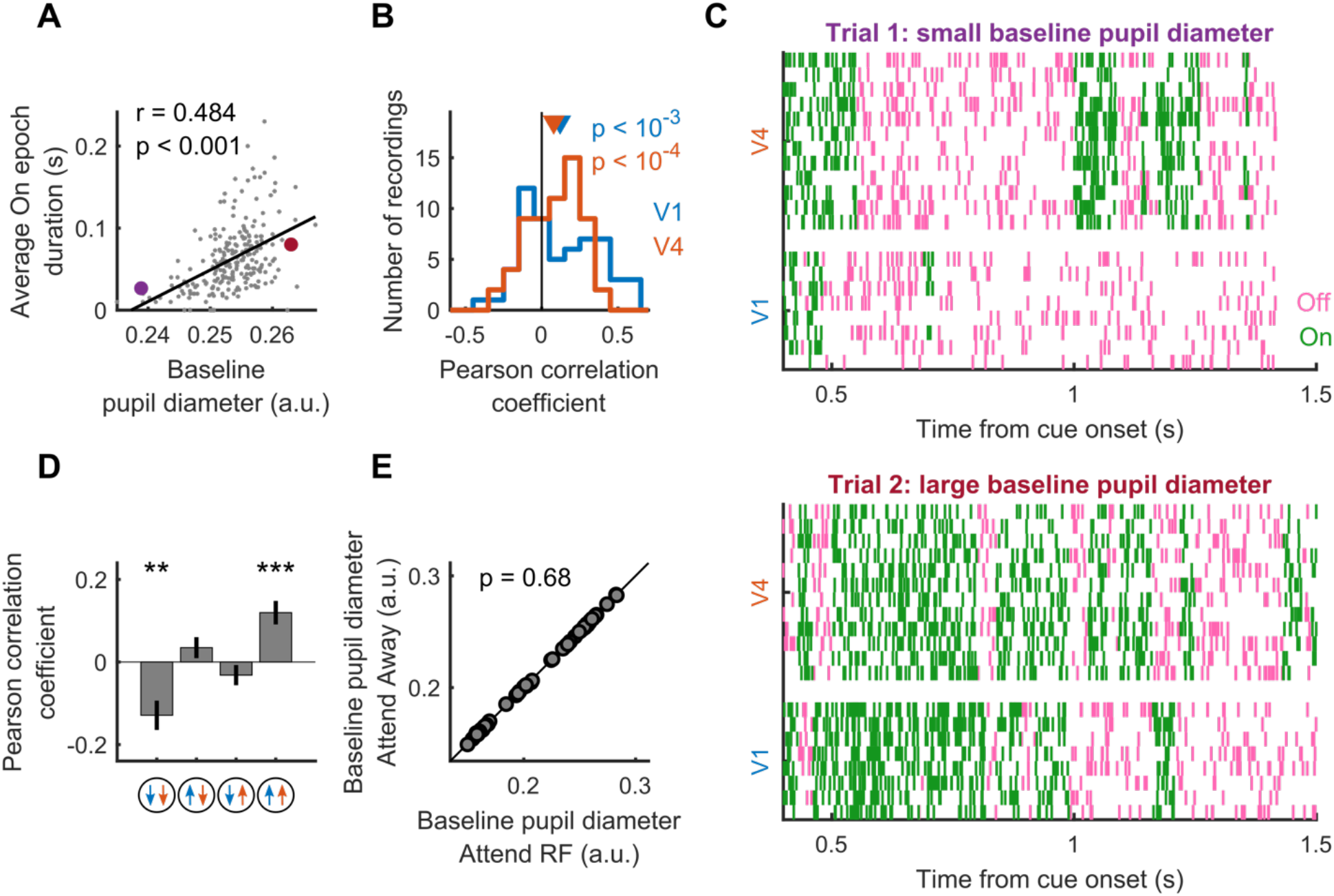
The relationship between baseline pupil diameter and On/Off episode durations. (**A**) Example recording showing that baseline pupil diameter is positively correlated to the average On episode duration in V1. Each dot represents one single trial, r is the Pearson correlation coefficient. The purple and red dot indicate the example trials used in panel **C**. (**B**) Across recordings, the average duration of On epochs in both V1 and V4 is positively correlated with the size of the baseline pupil diameter. (**C**) Two example trials in which the average On epoch duration is larger on the trial with larger (bottom) compared to the trial with smaller (top) baseline pupil diameter. (**D**) Across recordings, baseline pupil diameter is negatively (positively) correlated with the average epoch duration when both V1 and V4 are in an Off (On) phase. (**E**) The average baseline pupil diameter during attend RF conditions plotted against attend away conditions. There is no difference between attention conditions. Each dot represents a recording session. Statistics: Wilcoxon signed rank test (FDR corrected) (**B**, **D**, **E**) and Pearson correlation (**A**). Error bars and shaded regions denote ±1 SEM, and *, ** and *** indicate significance levels of p < 0.05, p < 0.01 and p < 0.001 respectively.

**Table S1.**
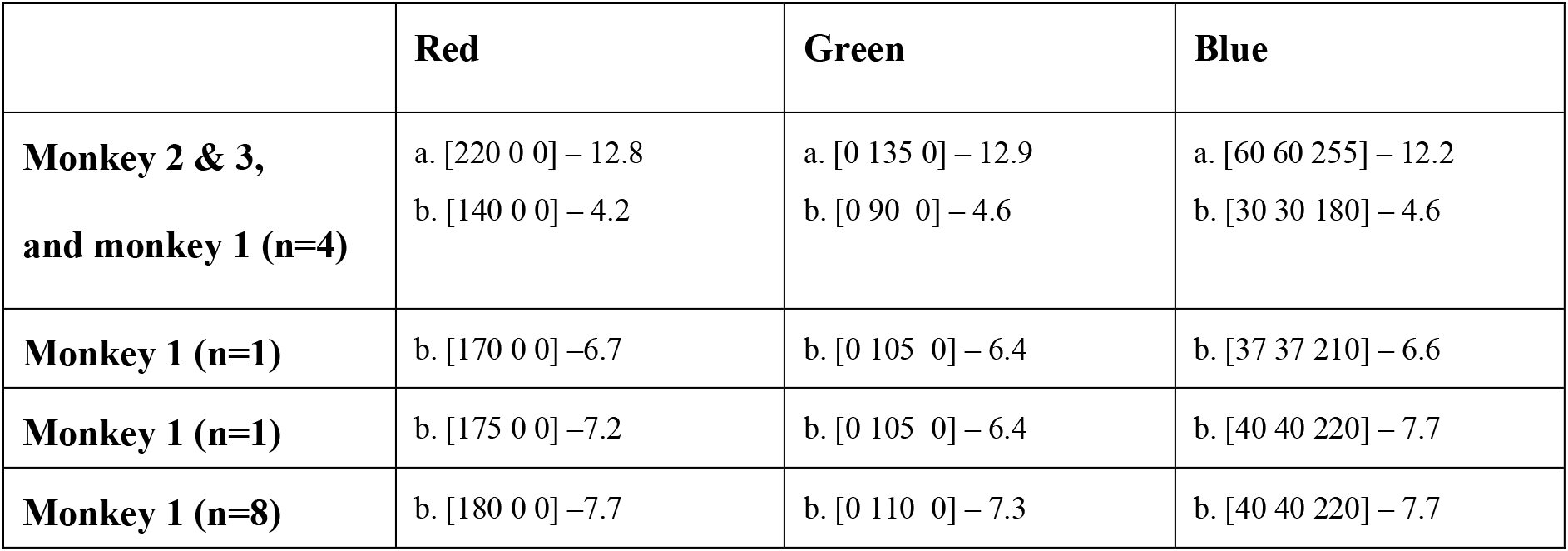
Color values used for the 3 colored gratings across recording sessions and subjects, indicated as [RGB] – luminance (cd/m^2^). a = Undimmed values, b = Dimmed values. For monkey 1, we used a variety of dimmed values across recordings.

|  | Red | Green | Blue |
| --- | --- | --- | --- |
| **Monkey 2 & 3,**<br>**and monkey 1 (n=4)** | a. [220 0 0] – 12.8<br>b. [140 0 0] – 4.2 | a. [0 135 0] – 12.9<br>b. [0 90 0] – 4.6 | a. [60 60 255] – 12.2<br>b. [30 30 180] – 4.6 |
| **Monkey 1 (n=1)** | b. [170 0 0] – 6.7 | b. [0 105 0] – 6.4 | b. [37 37 210] – 6.6 |
| **Monkey 1 (n=1)** | b. [175 0 0] – 7.2 | b. [0 105 0] – 6.4 | b. [40 40 220] – 7.7 |
| **Monkey 1 (n=8)** | b. [180 0 0] – 7.7 | b. [0 110 0] – 7.3 | b. [40 40 220] – 7.7 |

